# Lifespan Development of EEG Alpha and Aperiodic Component Sources is Shaped by the Connectome and Axonal Delays

**DOI:** 10.1101/2025.02.21.639413

**Authors:** Ronaldo Garcia Reyes, Ariosky Areces Gonzalez, Ying Wang, Yu Jin, Maria Luisa Bringas-Vega, Mitchell Valdes-Sosa, Cheng Luo, Peng Xu, Viktor Jirsa, Dezhong Yao, Ludovico Minati, Pedro A. Valdes–Sosa

**Affiliations:** Clinical Hospital of Chengdu Brain Science Institute, University of Electronic Science and Technology of China, 610054, Chengdu, China; Neuroinformatics, Cuban Neurosciences Center, 11300, Havana, Cuba; Center for Mind/Brain Science (CIMeC), University of Trento, 38123, Trento, Italy; School of Technical Sciences, University “Hermanos Saiz Montes de Oca” of Pinar del Río, Pinar del Rio, Cuba; China-Cuba Belt and Road Joint Laboratory on Neurotechnology and Brain-Apparatus Communication, University of Electronic Science and Technology of China, Chengdu, P. R. China; Laboratory for Brain Science and Artificial Intelligence, Southwest University of Science and Technology, Mianyang 621010, China; Aix Marseille Université, Institut National de la Santé et de la Recherche Médicale, Institut de Neurosciences des Systémes (INS) UMR1106; Marseille 13005, France

**Keywords:** Lifespan, Spectral Components, EEG-Dataset, Conduction Delays, Source Analysis, Alpha-Rhythms

## Abstract

We introduce *ξ* − *α*Net, a mesoscale generative model of cortical activity that models the EEG aperiodic (*ξ*) and *α*-rhythm (*α*) components via Hida-Matérn processes constrained by anatomical connectivity and interareal delays. This framework integrates fractional differential equations (modeling damping/resonance), decomposition of Spectral Granger Causality, and lifespan quantification. Bayesian inversion of the model, on cross-spectral rsEEG data from 1,965 participants aged 5-100 years (HarMNqEEG dataset), allows us to estimate cortical activity and effective connectivity patterns at 8,003-voxel resolution. *ξ* processes showed extended cortical networks, with occurrence probability increasing with age but amplitude peaking midlife (inverted-U trajectory). *α* processes exhibited a localization in the posterior areas of the cortex and two specialized networks (parieto-occipital, sensorimotor), with increasing localization probability and peak amplitude/frequency showing age-linked inverted-U trajectory. The model uniquely estimates global conduction delays, which are negatively correlated with *α* frequency and independent myelin concentration profiles, mechanistically linking neural propagation times to *α* regulation.

## INTRODUCTION

Spectral Components Models (SCM) have become a standard framework for decomposing neural oscillations into meaningful constituents. In [Tab. 1], we show a systematic review of the main SCM in the literature. In this paper, we analyze neural oscillations using the cross-spectrum, defined from an EEG vector time series 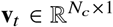:

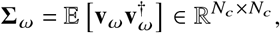

where *N*_*c*_ is the number of sensors, 𝔼 its the expectation over time trials, **v**_*ω*_ represents the Fourier transform of the EEG signal, and ***Σ***_*ω*_ denotes the cross-spectrum at frequency *ω*. The diagonal elements ***Σ***_*ω,ii*_ capture power spectra of individual channels, while the off-diagonal elements reveal channel interactions.

**Table 1.** Comparison of the main Spectral Components Models (SCM). All SCM approximate the Aperiodic Component as a Matérn process or its limiting cases.

| Model | Components Distribution |  |  | Scale | Cross | Non-Stationary | Anat-Priors | ESI | Connectivity | Ref. |
| --- | --- | --- | --- | --- | --- | --- | --- | --- | --- | --- |
| | AC | α-Peak | Others | | | | | | | |
| Zetterberg | AR(1) | AR(2) | — | 1 | — | — | — | — | — | [1] |
| $\xi - \alpha$ | $\mathcal{M}$ | $\mathcal{HM}$ | — | 1 | — | — | — | — | — | — |
| Multi $\xi - \alpha$ | $\mathcal{M}$ | $\mathcal{HM}$ | — | 1 | ✓ | — | — | — | — | [2] |
| Dipolar $\xi - \alpha$ | $\mathcal{M}$ | $\mathcal{HM}$ | — | 1 | ✓ | — | — | Dipolar | Functional | [3] |
| BOSC | $\mathcal{M}$ | $\chi^2$ | $\chi^2$ | log-log | — | — | — | — | — | [4] |
| IRASA | $\mathcal{M}$ | — | — | log-log | — | ✓ | — | — | — | [5] |
| FOOOF | $\mathcal{M}$ | $\mathcal{N}$ | $\mathcal{N}$ | log | — | ✓ | — | MNE | — | [6] |
| SPRINT | $\mathcal{M}$ | $\mathcal{N}$ | $\mathcal{N}$ | log | — | ✓ | — | — | — | [7] |
| PAPTO | $\mathcal{M}$ | $\mathcal{N}$ | $\mathcal{N}$ | log | — | ✓ | — | — | — | [8] |
| $\xi - \pi$ | Monotonic | Unimodal | Unimodal | 1 | — | — | — | — | — | [9] |
| Cortical $\xi - \alpha$ | $\mathcal{M}$ | $\mathcal{HM}$ | — | 1 | ✓ | — | — | eLORETA | Functional | [10] |
| $\xi - \alpha$NET | MVAR<br>+ $\mathcal{M}$ | MVAR<br>+ $\mathcal{HM}$ | — | 1 | ✓ | — | ✓ | Bayesian | Effective | — |

**Note:** **Dist.:** statistical distribution assumed for each component — AR: Autoregressive model; $\mathcal{M}$ : Matérn process; $\mathcal{HM}$ : Hida–Matérn process; $\mathcal{N}$ : Gaussian (normal); $\chi^2$ : chi-squared; Monotonic: constrained monotonic shape; Unimodal: single-peak shape. **Scale:** frequency axis scaling — 1: linear; log: logarithmic; log-log: log-power vs. log-frequency. **Cross:** multivariate modeling of the cross-spectra. **Non-Stat.:** non-stationary modeling capability. **Anat-Priors:** use of structural priors like Connectome + Delays. **ESI:** type of electrophysiological source imaging used (e.g., MNE, eLORETA, Bayesian). **Connectivity:** Functional: undirected association; Effective: directed/causal influence models.

Across the literature, two key Spectral Components (SC) consistently emerge [Fig. 1]: the Aperiodic Component (*ξ*-process), characterized by a monotonic decay with frequency, and the Periodic Component with a notable resonant peak in the *α*-band, 7–13 Hz), known as the *α*-Process [2,11–13]. The *ξ*-process is sometimes approximated by a 1 / *ω*^*β*^ decay, representing a fractional Brownian process in the time domain.

**Figure 1.**
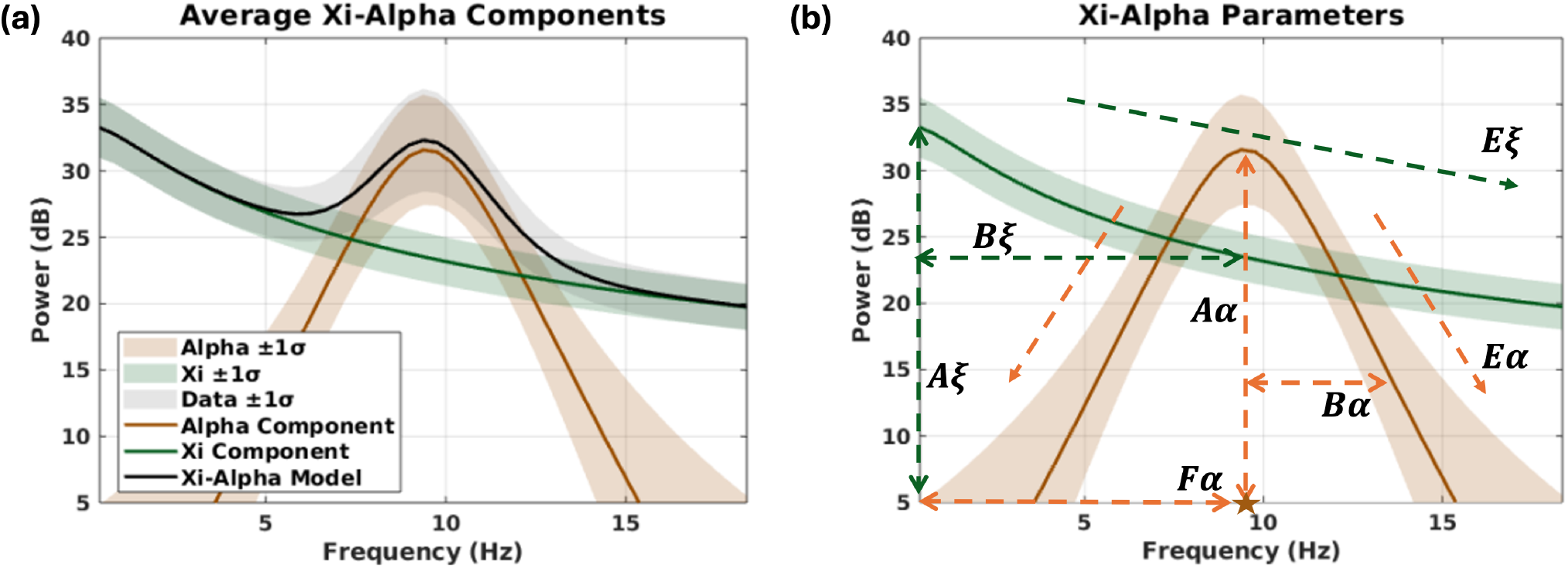
Xi–Alpha spectral decomposition for a representative subject. **(a)** The empirical EEG power spectrum (gray band) is modeled as a linear superposition of two physiologically interpretable and independent components with generalized Lorentz spectral profiles. The first component, the aperiodic process *ξ* (green line), captures the characteristic 1/*ω* spectral decay and reflects scale-free neural activity. The second component, the *α* -Process (orange line), models narrowband oscillatory activity centered within the 7–13 Hz range, corresponding to *α* -rhythms. The combined *ξ*–*α* model (black line) accurately captures the empirical spectral shape by summing the independent contributions of both components. Shaded regions indicate ±1 standard deviation across all EEG sensors of one subject. **(b)** The components are modeled at each sensor *i* using with a generalized Lorentz spectral distribution: (*ψ* (*ω* | *A, B, E*, *F*) = *A*/ 1 + *B* (*ω* − *F*)^2^ )^*E*^. where *A* is the amplitude (in dB), *B* is the bandwidth (in sec^2^), *E* is the spectral exponent, and *F* is the frequency shift (in Hz). The Aperiodic Component is given by *ξ*_*ω,i*_ = *ψ* (*ω* | *Aξ*_*i*_, *Bξ*_*i*_, *E ξ*_*i*_, 0), and the *α* spectral peak by *α*_*ω,i*_ = (*ψ* (*ω* | *Aα*_*i*_, *Bα*_*i*_, *E α*_*i*_, *F α*_*i*_ ) + *ψ* (*ω* | *Aα*_*i*_, *Bα*_*i*_, *E α*_*i*_, −*F α*_*i*_ ) )/2. Here, *Aξ*_*i*_ and *Aα*_*i*_ quantify the amplitude (i.e., the strength) of each component; *Bξ*_*i*_ and *Bα*_*i*_ control spectral width; *E ξ*_*i*_ and *E α*_*i*_ determine spectral decay; and *F α*_*i*_, shown by the orange asterisk on the frequency axis, denotes the Peak *α* Frequency (PAF) and characterize the modal frequency of the *α* -rhythms. For notational purposes, we will use the following notations: **A*ξ*, B*ξ*, E*ξ*, A*α*, B*α*, E*α*** and **F*α***. These vectors contain all the respective spectral parameters, with their dimensions corresponding to the number of channels or voxels, depending on the analysis [14].

**Figure 2.**
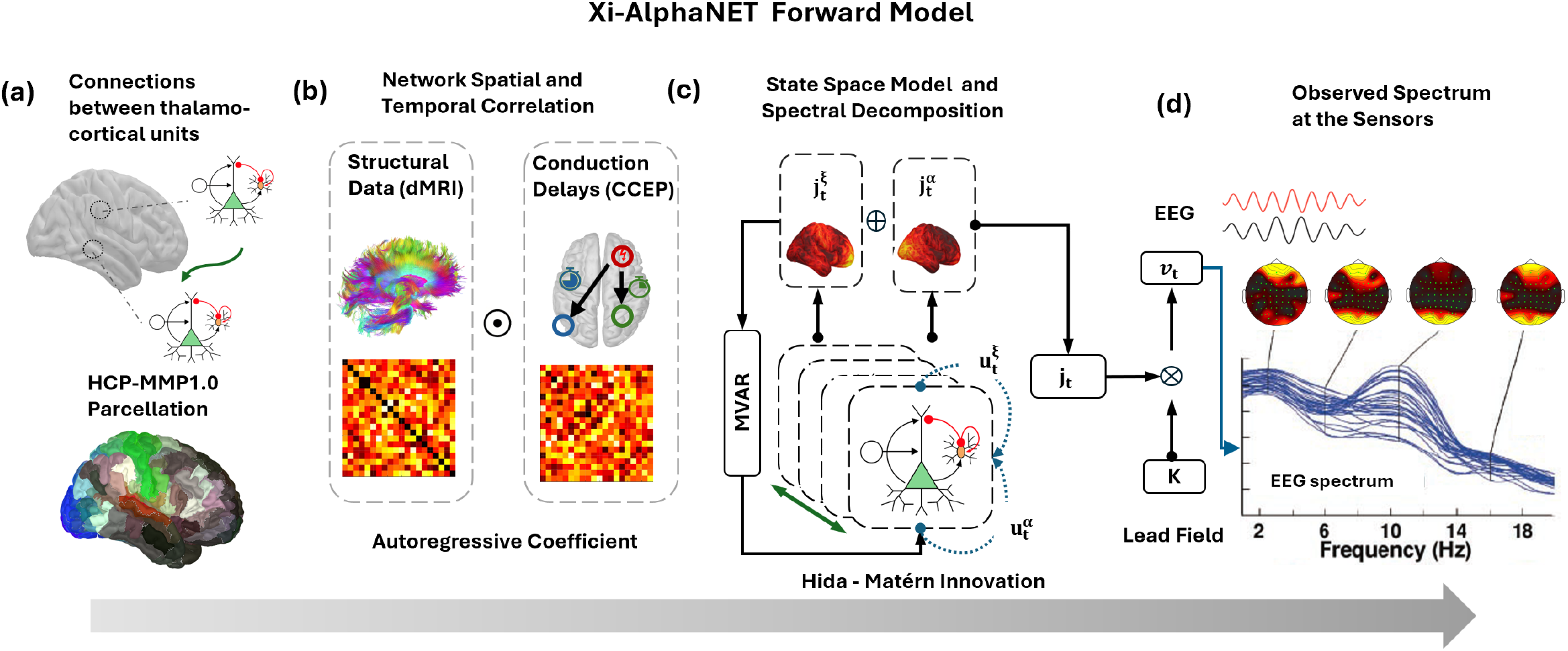
*ξ* − *α* NET Forward Model. (**a**) Connectivity between cortical generators is constrained with structural data from diffusion MRI (dMRI) tractography and conduction delays derived from cortico-cortical evoked potentials (CCEP), where HCP-MMP1 parcellation [21] is used to define the cortical generators. (**b**) Structural connectivity and conduction delay estimates shape the baseline spatiotemporal correlations of the cortical network. (**c**) The cortical activity (**j**_*t*_ ) arises from the linear superposition of two independent spectral processes, where 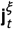 accounts for the aperiodic component and 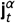 for the *α* spectral peak. Each activation is modeled using an independent MVAR state-space model with colored Hida-Matern innovations 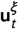 and 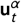. The autoregressive coefficient of the MVAR model is shaped by the connectome and the conduction delays and allows for modeling non-local interactions between cortical generators, while the independent Hida Matern innovations represent local processes at each thalamocortical unit that allows for introducing Lorenzian spectral profiles for each spectral process, which parameters have sparse distribution on the cortical surface. (**d**) Cortical activity is then projected to the sensor using the lead field matrix (**K**), which generates the observed EEG spectra in concordance with the electrophysiology. This framework enables the estimation of cortical SC, conduction delays, and source effective connectivity from EEG/MEG recordings, supporting large-scale normative studies of cortical maturation. The symbol ⊙ denotes the Hadamard (element-wise) product used to introduce a spatiotemporal decomposition of the interactions between generators of neural activity, ⊕ denotes the sum, and ⊗ denotes the standard matrix product.

Two complementary approaches have historically shaped SCM development [Tab. 1]. The foundational work of Zetterberg et al. [1] adopted a time domain approach using Autoregressive Moving Average (ARMA) models in the time domain to estimate EEG spectra. In contrast, Pascual-Marqui et al. [2] pioneered a frequency domain approach with the *ξ* − *α* model, representing EEG spectra as a mixture of independent stochastic processes: the *ξ* and *α*-processes. Pascual’s adoption of a generalized Lorentzian profile—originally inspired by spectroscopy—enabled efficient fitting to EEG spectra [Fig. 1]. Subsequent work by Amador et al. [14] shows the success of this model in capturing developmental changes. As we are going to show in this paper, the Lorentzian spectral profiles that characterize the *ξ* − *α* model naturally arise from time-domain processes governed by the Hida–Matérn class. This connection extends beyond the aperiodic component, suggesting that all EEG components are driven by fractional stochastic dynamics, a viewpoint missing in contemporary SCM.

Recent SCM have diversified, adopting the frequency-domain framework introduced by Pascual through the *ξ*–*α* model [Tab. 1]. Univariate SCM such as BOSC [4], IRASA [5], FOOOF [13], SPRiNT [7], PAPTO [8], and the recent *ξ*–*π* model [9] offer a range of parametric and non-parametric teqniques.

On the other hand, we also have a multivariate approach to model the off-diagonal elements of the cross-spectra. An early example it is the multivariate *ξ*-*α* model [2], though restricted to sensor-level EEG. Valdés-Sosa et al. [3] later proposed the first source-level formulation, modeling *ξ* as an isotropic process on the cortical surface and *α*-rhythms as two correlated stochastic dipole processes with fixed orientations and magnitudes. More recently, Pascual et al. [10] introduce Cortical *ξ*–*α*, which extends the *ξ*–*α* model to cortical source space using eLORETA for spectral functional connectivity estimation.

As we conclude from [Tab. 1], current approaches to estimating the neural sources of SC typically follow a two-step strategy: (i) localizing activity to putative source sensors, followed by (ii) fitting SCM parameters. All these techniques neglect key anatomical and functional constraints, such as structural connectivity, axonal delays, and conduction delays, that shape the spatiotemporal propagation of oscillatory activity [15,16]. As we have previously shown, such a two-step strategy can lead to biased estimates due to misspecified source covariance matrices [17].

Solutions to all these challenges can be found within large-scale initiatives such as the Digital Twin Brain and The Virtual Brain Project [18–20], which integrate rich neuroanatomical data to capture neural oscillations more realistically. However, they require substantial computational resources, which limits their direct practical applicability in large-scale normative studies. To overcome these limitations, here we introduce *ξ* −*α*NET, a mesoscale generative model of cortical activity formulated as a sparse, structurally constrained network of Hida–Matérn processes. Unlike previous SCM approaches, *ξ* − *α*NET allows the joint inference of source-level spectral dynamics and inter-regional effective connectivity. Using Bayesian inversion, we achieve both computational efficiency and high spatial resolution—demonstrated at 8,003 cortical voxels—on the HarMNqEEG dataset, comprising 1,965 participants across 9 countries and including resting-state EEG (rsEEG) cross-spectrum data from multiple acquisition devices. This framework allows the construction of large-scale, normative research on developmental trajectories of cortical dynamics.

## RESULTS

### Xi-AlphaNET as a Sparse Structurally Constrained Network of Hida-Matérn Process

#### Forward Model

We introduce *ξ*–*α*NET, a mesoscale generative model of cortical dynamics that accounts for EEG observations by integrating voxel-wise spectral processes with anatomically grounded structural connectivity and conduction delays. To start, the observed EEG time series 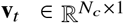 is modeled as a linear projection of distributed cortical activity:

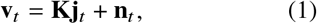

where 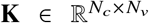 is the lead-field matrix and **n**_*t*_ is zero-mean gaussian noise. We aim to model the time-domain cortical activity 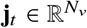 at each thalamocortical unit under three constraints: (i) the model must incorporate structural priors, reflecting structural connectivity and conduction delays across neurotracts, that shape the spatiotemporal correlations in a causal way; (ii) the spectral dynamics must arise as a linear superposition of two statistically independent components, corresponding to aperiodic and oscillatory generators; (iii) and the resulting power spectrum must exhibit Lorentzian profiles, consistent with empirical spectra observed across multiple frequency bands in EEG/MEG recordings.

In order to satisfy the first condition (i), we model the cortical dynamics within a structurally constrained Multivariate Autoregressive (MVAR) network, where the autoregressive coefficients are informed by the structural priors [22,23]. However, simple MVAR or AR models fail to reproduce the full characteristics of Lorentzian spectral profiles of neural oscillations, particularly in the *α*-band spectral properties. Therefore, it is necessary to use MVAR with colored innovations.

The second condition (ii) can be warranted if we assume that the cortical activity **j**_*t*_ is the linear superposition of two independent stochastic processes that model the activations of the independent SC. This approach is aligned with the linear decomposition of the cross-spectrum in SC used on the Multivariate *ξ*–*α* proposed by Pascual et al. [2,3]. However, despite the success of *ξ*–*α* models, the underlying time-domain mechanisms capable of generating Lorentzian spectral profiles have never been explained. Here, we report an advance that addresses this gap: *Lorentzian spectral profiles can be generated by real-valued fractional stochastic processes belonging to the Matérn class*. These are Gaussian processes with a temporal delayed structure or autocorrelation. Matérn processes have found extensive use in spatial statistics and machine learning. However, the application of this process to model cortical time series has been overlooked within the neuroscience community. As shown in [Tab. 1], we found that the aperiodic component of neural oscillation can be explained in terms Matérn process, which provides a principled time-domain origin of the *ξ*-process defined by Pascual et al. [2,3].

Resonant spectral peaks, such as those observed in the *α*-band or other bands, present a significant challenge because they reflect intrinsic oscillatory dynamics that conventional Matérn processes, by themselves self cannot fully capture. Early work, like the thalamocortical Robinson model [24,25], shows that damped differential equations can generate oscillatory modes through the interplay of damping and delays. However, the Robinson model cannot reproduce the full range of spectral patterns observed in the *α*-band, particularly the Lorentzian spectral profiles with arbitrary exponents, which govern the smoothness of the corresponding time-domain processes introduced by Pascual et al. [2,3,10]. Properly modeling these features requires the use of fractional stochastic differential equations, as implemented in oscillatory Matérn processes [26].

Although oscillatory Matérn processes introduce a resonant spectral peak, the time domain requires complex-valued stochastic processes [26]. This is inconsistent with the real-valued nature of neurophysiological signals. Therefore, in order to address this limitation, we model the *α*-rhythm within each thalamocortical unit using the Hida-Matérn process. Unlike the oscillatory Matérn process, the Hida-Matérn formulation yields an analytically tractable, physically interpretable, and real-valued process in a time-domain model that aligns with the expected emergent properties of thalamocortical activity [27]. Importantly, Hida-Matérn processes are exact solutions to fractional stochastic differential equations. This mathematical framework has been successfully applied to complex phenomena such as turbulence and anomalous diffusion, and now, through our model, to neural oscillations.

Therefore, one of the possible linear models of cortical activity that satisfies all these three conditions can be represented using an MVAR model with colored innovations as follows:

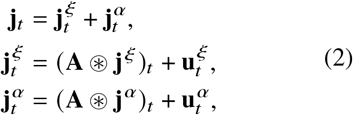

where 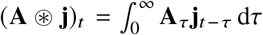 denotes delayed convolution over interactions. The kernel 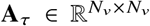 is a delay-dependent matrix of autoregressive coefficients with a null diagonal that governs non-local spatiotemporal propagation between thalamocortical generators. As previously demonstrated by our group, the concatenation of these delay-dependent interactions defines what we term the Delayed Connectome Tensor [28,29]. This tensor is influenced by the connectome and the conduction delays. Therefore, to account for structural prior information, we opt for an embedded approach to model the effects of anatomical structure by parametrizing the convolution kernel using a spatiotemporal decomposition of interactions via Hadamard product [23,28]:

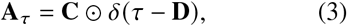

where *δ*(·) is the Dirac delta function applied element-wise, and ⊙ denotes the Hadamard (element-wise) product. The matrices 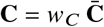 and 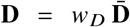 encode subject-specific coupling strengths and conduction delays, scaled from population-level priors 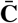 and 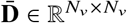. Where *w*_*D*_ and *w*_*C*_ are structural modulatory weights that are selected within the search space that warrant that the cortical activity represents a weakly stationary process and physiologically plausible values of conduction delays, respectively. This formulation links anatomical connectivity to dynamical interactions, in line with the framework proposed by Fukushima et al. [22]. The driven colored Gaussian noise process 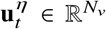, with *η* ∈ {*ξ, α*}, models the within thalamocortical unit dynamics for each spectral component as a collection of voxel-wise independent Hida-Matérn processes. Each process is zero-mean and temporally stationary, with auto-covariance structure given by:

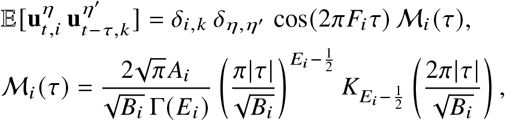

where *δ*_*i,k*_ and 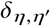 are Kronecker deltas enforcing independence across voxels and SC, respectively (not to be confused with the Dirac delta function *δ* (*τ*) used to describe autoregressive coefficients in the convolutional model). The function ℳ_*i*_ (*τ*) denotes the Matérn kernel shaping the temporal envelope. Here, *A*_*i*_ > 0 is the amplitude, *B*_*i*_ > 0 controls the temporal bandwidth (inversely related to peak sharpness), 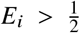 determines the smoothness, and *F*_*i*_ ≥ 0 specifies the central frequency of oscillation. *K*_*v*_ (·) denotes the modified Bessel function of the second kind, and Γ(·) is the Euler Gamma Function [27].

As a result of this formulation, the corresponding power spectral densities of the aperiodic and *α* components at voxel *i*, 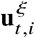 and 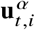 coincide with the spectrum parameterization ***ξ*** _*ω,i*_ and ***α***_*ω,i*_ introduced by Pascual et al. [2,10].

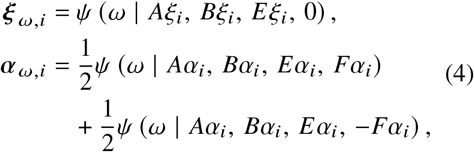

where the function *ψ* (*ω* | *A, B, E, F*) denotes a generalized Lorentzian spectral peak centered at frequency *F*, with amplitude *A*, bandwidth parameter *B*, and spectral exponent *E*. These parameters fully determine the shape and scale of each generator’s spectral profile [see Fig. 1]. Throughout this paper, we are going to refer to these quantities as the spectral parameters of the model. In order to enforce sparsity over the SC, we apply a Group Lasso regularization across the parameters of each cortical generator. This is done in order to ensure that if a given cortical generator does not contribute to a particular spectral component power (e.g., *α* or aperiodic), then all its corresponding spectral parameters are simultaneously driven to zero at the cortical generator [30].

As a consequence of this formulation, *the generative model of the ξ-αNET represents a sparse, structurally constrained network of Hida-Matérn processes in the time domain*, determined by two classes of parameters: (i) the spectral parameters 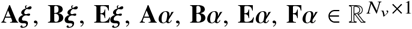, which shape the aperiodic and *α* spectral profiles at each voxel; and (ii) the structural parameters *w*_*C*_ and *w*_*D*_, which define the spatiotemporal dynamics of the thalamo-thalamo-cortical network. Full mathematical derivations and formal model details are provided in Supplementary Materials A & B.

#### Spectral Effective Connectivity

Within the *ξ* − *α*NET framework, the cortical activity of each spectral component is modeled by its own MVAR process with Hida-Matérn innovations. This formulation implies that each spectral process defines an independent causal structure between cortical generators. Consequently, *spectral decomposition naturally entails a decomposition of effective connectivity*, in line with the decomposition of functional connectivity obtained using lagged coherence by Pascual et al. [10].

However, mapping effective connectivity within the *ξ* − *α*NET model cannot be done straightforwardly using standard measures such as Isolated Effective Coherence (iCoh), Lagged Coherence, Spectral Granger Causality (SGC), Partial Directed Coherence (PDC), or Noise Contribution Ratio (NCR) [16,31–33]. These connectivity measures assume temporally uncorrelated (white) innovations within the generative model. Therefore, in order to use them to map effective connectivity, it is necessary to express them in terms of the System Transfer Function (TF) after whitening [31,33]. In the case of the *ξ* − *α*NET model, each spectral component yields a frequency-resolved transfer function with explicit parametric expressions:

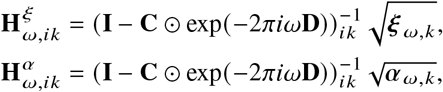

where **A**_*ω*_ = **C** ⊙ exp (−2*πiω***D**) denotes the frequency-resolved autoregressive coefficient matrix, incorporating structural connectivity and conduction delays. From these expressions, several key conclusions emerge. First, for directed propagation to occur from the source generator *k* into a target generator *i*, the spectral process must be active at the source generator (i.e., ***ξ*** _*ω,k*_ or ***α***_*ω,k*_ is nonzero) and also an anatomical path must exist between the source and target generators (i.e., **C**_*ik*_ ≠ 0). If ***ξ*** _*ω,k*_ or ***α***_*ω,k*_ → 0, then propagation is effectively sup-pressed. Second, each spectral process also defines a distinct cross-spectrum:

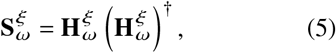

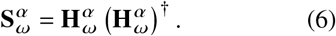

From this cross-spectra, we can compute the SGC to approximate the effective connectivity, as shown by Friston et al., [16,32,33] for each spectral process. The reason that motives the use of SGC and not another measure of connectivity like iCoh and PDC previously mentioned is that when you manipulate analytically these expressions, they introduce spurious correlations due to vanishing innovations. The way to properly introduce these other measures of connectivity to handle the influence of indirect pathways will be the focus of further investigation.

Finally, by manipulating the cross-spectra of each spectral process, we derive an expression for the spectral power density at each generator:

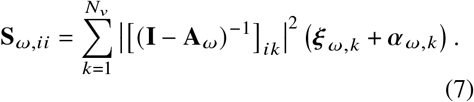

Therefore, within the *ξ* − *α*NET framework, spectral power emerges as a consequence of network propagation, with the Green function **I** ( − **A**_*ω*_) ^−1^ characterizing the redistribution of power across the network. In the limit of absent inter-generator coupling (**A**_*ω*_ → 0), the model reduces to a purely local formulation, recovering the classical voxel-wise *ξ* − *α* model applied to the sources [2,3].

#### Bayesian Model Inversion

We estimate the parameters of *ξ*–*α*NET from rsEEG cross-spectral data in the HarMNqEEG dataset [36]. For each subject *j*, the model input is the empirical cross-spectrum 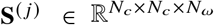, which represent the frequency-resolved sensor covariance. Parameter estimation is framed as a Maximum A Posteriori (MAP) problem, combining the likelihood of the model-predicted source spectrum with priors on spectral and structural parameters (see Supplementary Materials C).

The resulting MAP objective function is non-smooth and non-convex, so we adopt a two-step profile likelihood strategy. First, the structural parameters for the connectivity and delay are estimated using Bayesian optimization, constrained by physiological priors from lifespan studies [35]. This defines subject-specific structural anchors. Second, spectral parameters are inferred using Stochastic FISTA with Nesterov acceleration [37], initialized from 50 random seeds within bounded domains. Model complexity is penalized using BIC, and regularization hyperparameters are tuned per subject via Bayesian optimization (see Supplementary Materials D–F).

### One-step *ξ*–*α*NET approach outperforms two-step pipelines in spectral components reconstruction

We benchmarked *ξ*–*α*NET, simulating physiologically grounded synthetic EEG data and compared it against two standard SCM pipelines. To this end, we performed 100 simulations of a biophysically informed neural mass model, formulated as a second-order differential equation with a similar structure to the Jansen–Rit model [28]. A neural mass was assigned to each of the 360 regions in the HCP-MMP1 parcellation [21]. Within each neural mass, the cortical activ-ity is modeled as a linear superposition of two non-independent damped resonators: one aperiodic (0 Hz) and one within the *α*-band (7–13 Hz). Nonlinear interactions are described using a sigmoidal function on delayed interregional inputs, enabling physiologically grounded cross-frequency coupling.

When we want to benchmark through simulations an inverse solution method like the *ξ* − *α*NET, it’s important to avoid as much as possible what is known as inverse solution crime [38], which consists of using the same generative model to generate synthetic data and then validating. In the case of this simulation, the generative model defined to generate the synthetic data is different from the one that defines the *ξ*–*α*NET, which is based on a linear MVAR with colored innovations, which minimizes the risk of inverse solution crime. To introduce variability and promote topological diversity of the *α*-rhythms, each simulation was initialized with different random conditions.

Inter-regional coupling was constrained using the structural connectivity atlas estimated by Rosen et al. [34] from diffusion MRI, while conduction delays used are from the F-TRACT atlas of cortico-cortical conduction delays based on CCEP recordings [35]. These structural priors specify the spatiotemporal evolution of cortical activity in a physiologically plausible manner. The simulated source cross-spectrum was then projected to the sensors using the lead field matrix. In order to emulate empirical sensor noise, we added complex Wishart-distributed noise. A full description of the simulations its developed in the Supplementary Material H.

We assessed the ability of *ξ*–*α*NET to recover both the source power spectrum and functional connectivity, comparing its performance to two widely used two-step approaches: MNE followed by FOOOF [13], and eLORETA followed by *ξ*–*α* model. These methods represent standard practice in the field, where inverse modeling and spectral fitting are performed as separate steps. As shown in [Fig. 3(b)], *ξ*–*α*NET achieved substantially lower mean squared error in reconstructing the ground-truth source spectra, capturing both the shape and spatial distribution of the aperiodic and *α* components more precisely than the baselines.

**Figure 3.**
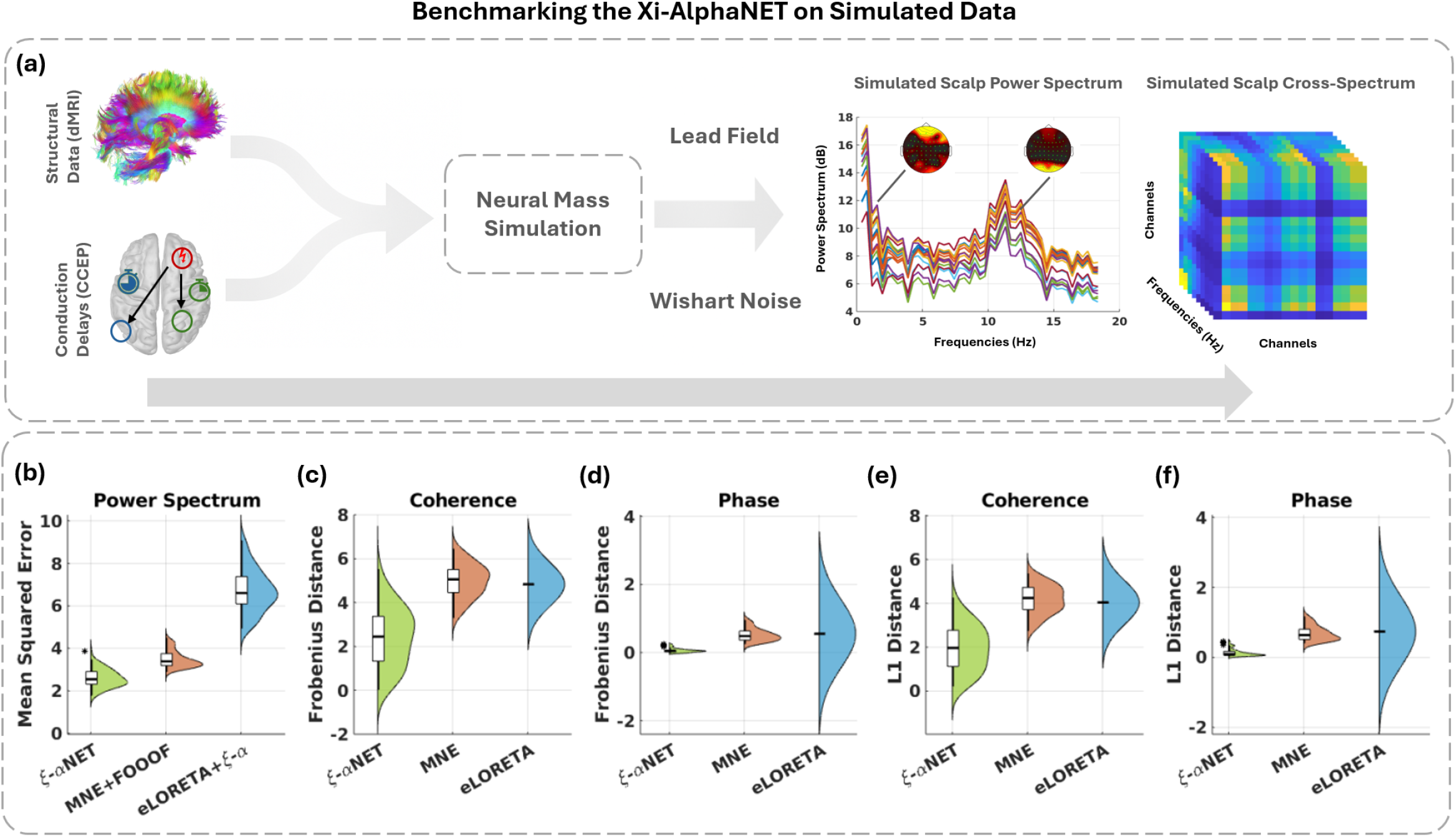
Validation of the *ξ*-*α* NET Model on synthetic EEG data (Nsim = 100). (**a**) To simulate synthetic EEG data, we use a Delayed Neural Mass Model with a different generative model from the one that defines the *ξ*-*α* NET Model and that incorporates the structural connectivity atlas estimated using dMRI by Rosen et al. [34], and the conduction delay atlas estimated using CCEP by Lemarechal et al. [35], both of them using the HCP-MMP1parcellation Glasser et al., [21] Then, after each simulation, the cortical time series in the 360 neural masses, we take the covariance between generators at each frequency bin to estimate the cortical cross-spectrum tensor. Then we map it into the sensor cross-spectrum tensors using the lead field and Wishart noise, which, as we depict, has an aperiodic component and a random peak within the *α* -band and a topographical distribution in concordance with electrophysiology. (**b-f**) Performance comparison of *ξ*-*α* NET against MNE+FOOOF and eLORETA+*ξ*-*α* for various metrics: power spectrum, coherence, phase with the Frobenius distance, and L1 distance. *ξ*-*α* NET demonstrates superior performance across all benchmarks, suggesting that one-step spectral estimation yields superior performance than two-step approaches.

To evaluate connectivity estimation, we computed coherence and phase-lag matrices and quantified reconstruction accuracy using Frobenius and L1 distances. These metrics respectively capture global mismatches and sparsity errors. Across all conditions—coherence [Fig. 3(c,e)] and phase [Fig. 3(d,f)]—*ξ*–*α*NET outperformed both MNE and eLORETA-based approaches, yielding tighter error distributions and smaller median distances. Its superior phase-lag estimates, in particular, reflect the benefit of explicitly modeling conduction delays within the inversion.

Overall, these results suggest that a one-step inversion framework such as *ξ*–*α*NET—which jointly estimates source activity, spectral parameters, and connectivity—yields more accurate reconstructions than conventional two-step pipelines. By embedding structural priors and spectral constraints directly into the generative model, *ξ*–*α*NET provides a robust and principled solution for EEG-based network inference.

### Spectral components exhibit distinct cortical localizations and effective connectivity patterns across the Lifespan

To map the distribution of SC across the Lifespan, we performed Bayesian model inversion of *ξ* − *α*NET on the entire HarMNqEEG dataset. For each subject *j*, the observed EEG cross-spectrum 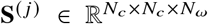 was used to estimate individualized source-level SC via MAP optimization. We then estimated the age-dependent probability of expression of each SC at each cortical or Probability Atlas by applying a Nadaraya-Watson kernel density estimator to the set of amplitudes {**A*ξ*** ^( *j*)^, **A*α***^( *j*)^ } _*j*_ across all subjects. This nonparametric regression approach enabled robust estimation of conditional probabilities over age, providing spatially resolved maps of SC expression trajectories throughout the lifespan [39]:

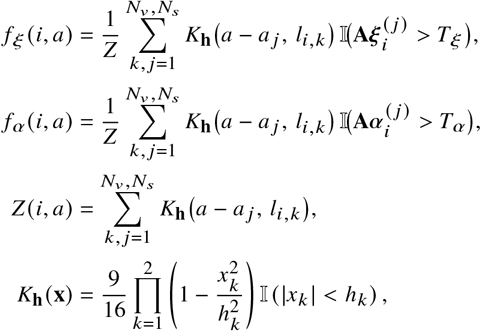

where *K*_**h**_(·) is the Epanechnikov kernel with a smoothing parameter **h** = (*h*_1_, *h*_2_) ^⊤^ selected by 10-fold cross-validation. Here, *l*_*i,k*_ is the distance between voxels *i* and *k, Z* (*i, a*) is a normalization constant, and 𝕀 (·) is an indicator function that equals one if the SC power exceeds a threshold (*T*_*ξ*_ or *T*_*α*_). These thresholds are set automatically by fitting a mixture of two Gaussian distributions on the power of the SC through the use of Expectation Maximization, and it is defined as the point at which the two Gaussian distributions overlap (see details in Supplementary Material G). From the density estimation, we compute the spatial distribution of SC as the marginal distribution across age groups.

On the other hand, in order to approximate the effective connectivity across the Lifespan, we first estimate the source cross-spectrum for each spectral component and subject in the HarM-NqEEG dataset using Eqs. (5,6). Then, we compute the average source cross-spectrum for each age group and spectral component. For that, it is necessary to take into account that, after regularization, the cross-spectrum is positive definite and, therefore, resides on the Rie-mannian manifold of positive-definite matrices, which possesses an intrinsic nonlinear structure. It is known that in these spaces, traditional linear estimators of central tendency as the Euclidean mean 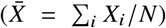, fail to account for the geometric structure of the cross-spectra, leading to spurious causal interactions due to poor statistical handling. In consequence, it is necessary to use more advanced methods, such as the Fréchet mean, under an appropriate metric. In this paper, we employ the Log-Euclidean metric 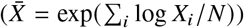 to compute the Fréchet mean, which is both computationally efficient and well-suited for analyzing large datasets like the HarMNqEEG. Further details regarding this methodology can be found in the works developed by our group [36,40,41]. After obtaining the mean cross-spectrum for each age group and spectral process, we map it into the corresponding SGC to approximate the effective connectivity.

The results of our analysis of the HarMN-qEEG dataset are presented in [Fig. 4], which depicts the Developmental Atlases of the Localization Probability and the SGC for both the *ξ* and *α*-processes. The probability atlas for the *α*-process shows a prominent localization in the posterior areas, with only modest topological variance across the Lifespan [Fig. 4(a,c)]. On the other hand, the effective connectivity network of the *α*-process reveals two distinct specialized networks: A parieto-occipital network related to the visual processing and a sensorimotor network capturing connectivity patterns of the *μ*-rhythms [Fig. 4(b)]. As can be depicted in [Fig. 4(d)], these two specialized networks are disconnected at the early stages of development, and the connectivity between the sensorimotor and the parieto-occipital network increases with age.

**Figure 4.**
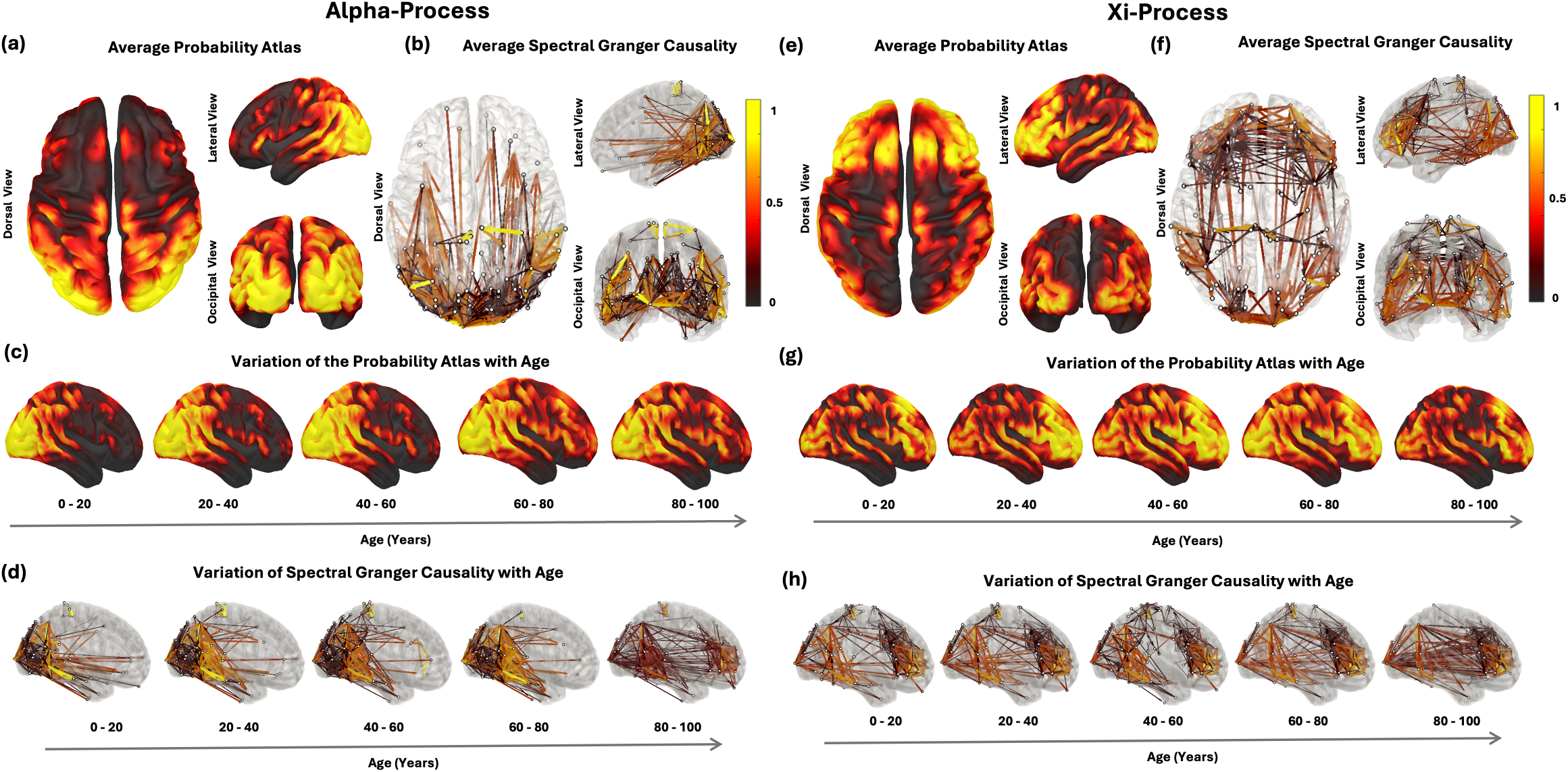
Lifespan Distribution of Probability Atlas and Spectral Granger Causality for each Spectral Component. Panels **(a)**–**(d)** show results for the *α* -Process (spectral peak in the *α* -band), and panels **(e)**–**(h)** show results for the *ξ*-Process (aperiodic component). All estimates are based on *ξ*-*α* NET model inversion using the HarMNqEEG dataset (*N* = 1965 participants, *N*_*v*_ = 8003 voxels). **(a**,**e)** Average Probability Atlas: For the *α* -Process, high probabilities localize in the posterior cortex with a gradient towards frontal regions, indicating spatial localization of *α* -rhythm generators. In contrast, the *ξ*-Process shows widespread distribution across the entire cortical surface, suggesting more diffuse generation. **(b**,**f)** Average Spectral Granger Causality (SGC): Computed from the Log-Euclidean mean of the cross-spectra and mapped to SGC, revealing that for the *α* -Process causal interactions are mainly within the posterior areas of the cortex, with some projections to frontal regions consistent with posterior-to-frontal propagation. The *α* -process also depicts two specialized networks, a parieto-occipital network related to visual processing and a sensorimotor network related to the *µ*-process. On the other hand, the *ξ*-Process has a widespread, bidirectional connectivity that emerges across multiple cortical regions. **(c**,**g)** Variation of the Probability Atlas with Age: The *α* -Process shows have low variability of cortical distribution, being predominantly localized in the posterior areas. While the *ξ*-Process remains widespread, but shows increasing probability in frontal regions with maturation. **(d**,**h)** Variation of SGC with Age: For the *α* -Process, the two specialized networks (parieto-occipital and sensorimotor) are disconnected at an early stage of development, and connectivity between them increases with age. On the other hand, the *ξ*-Process shows increasing bidirectional connectivity between posterior and frontal regions.

In contrast, the cortical generators of the *ξ*-process (aperiodic component) are distributed more widely across the cortical surface throughout the Lifespan [Fig. 4(e)]. Unlike the *α*-process, which displays a localized connectivity pattern, the *ξ*-process exhibits a widespread effective connectivity network, encompassing causal interactions across multiple cortical regions [Fig. 4(f)]. With aging, we appreciate an increased bidirectional connectivity between posterior and frontal regions [Fig. 4(g,h)].

### Spectral Components localization correlates with significant inverted U-Shape developmental trajectories and an increase of localization probability

The output of the *ξ − α*NET inversion yields, for each subject *j*, voxel-wise estimates of spectral amplitude for both the aperiodic component **A*ξ*** ^( *j*)^ and the *α*-band component **A*α***^(*j*)^, as well as the corresponding PAF map **F*α***^(*j*)^. While age-related changes in these features have been widely reported at the sensor level [36,42–44], their spatially resolved trajectories at the cortical source level have not been systematically characterized across the Lifespan.

Due to the Group Lasso prior regularization on the generative model of the *ξ*-*α*NET, the distributions of **A*ξ*** ^(*j*)^, **A*α***^(*j*)^, and **F*α***^(*j*)^ are inherently sparse, with many voxels exhibiting exact zeros. The resulting execs of zeros in the data its known as zero inflation, which violates key assumptions of conventional Gaussian-based regression. Therefore, in order to appropriately map the developmental trajectories of the SC, we employed a Zero-Inflated Gaussian (ZIG) regression model. The use of this type of regression model allows us to account for the sparsity, non-negativity, and nonlinear age effects, providing a robust statistical basis for mapping developmental trends of source-resolved spectral activity [45].

We employed a voxel-wise ZIG regression model of the form (Supplementary Materials I):

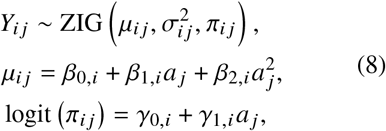

where *Y*_*ij*_ denotes the response variable—either **A*ξ*, A*α***, or **F*α***—in voxel *i* for subject *j*, and *a* _*j*_ is the subject’s age. On the other hand the function logit (*p*) = log (*p* / (1 − *p*) ) is the logit link, and *μ*_*ij*_ is the conditional mean (CM) which captures the expected value of the positive (non-zero) portion of the distribution.

[Fig. 5] summarizes the Developmental Atlas of spectral EEG components captured by the ZIG regression model applied to *ξ*–*α*NET main spectral parameters estimated from the HarMNqEEG dataset. Panels 5(a)–(c) show voxel-wise statistical maps of age effects on the zero-inflation probability (ZIP, top row) and the quadratic curvature of the conditional mean (CM, bottom row) for each of the three spectral variables. For the **A*ξ*** [Fig. 5(a)], we observe a widespread decrease in ZIP with age (i.e., *γ*_1_ < 0), indicating that aperiodic activity becomes increasingly detectable across the cortex in older individuals. At the same time, the CM exhibits significantly concave (inverted U-shaped) trajectories across nearly all cortical regions (*β*_2_ < 0), suggesting that **A*ξ*** increases in early life, peaks in mid-adulthood, and declines thereafter.

**Figure 5.**
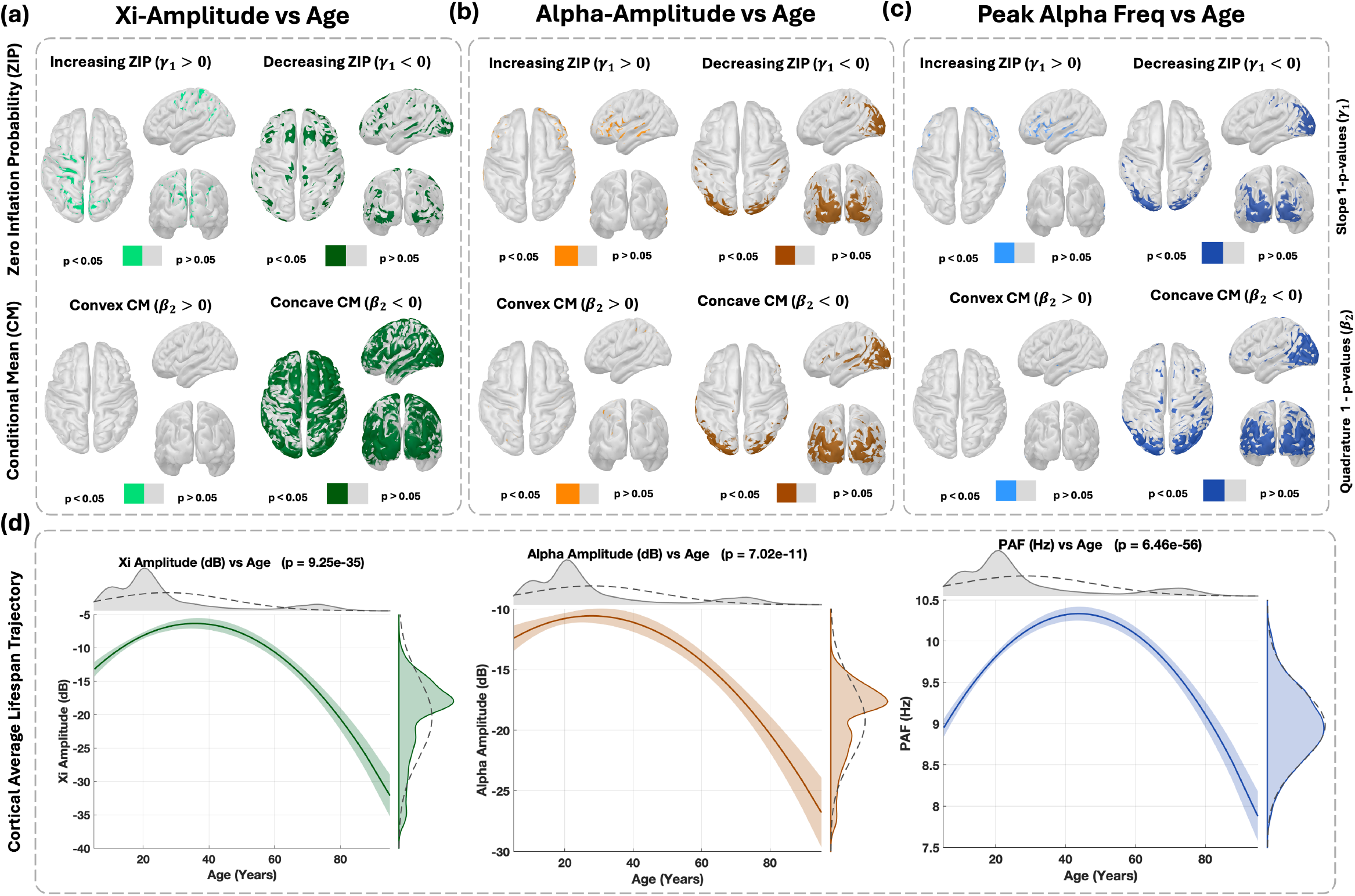
Developmental Atlas of spectral EEG components estimated using zero-inflated Gaussian (ZIG) regression on the HarMNqEEG dataset (*N* = 1965). We applied voxel-wise ZIG regression to estimate the developmental changes of the spectral parameters estimated from *ξ* − *α* NET model on the HarMNqEEG dataset. On each voxel, we jointly estimated the conditional mean (CM) of non-zero values and the probability of zero inflation (ZIP) as functions of age. For each component, spatial maps show voxel-wise distributions of 1 − *p* values associated with the linear ZIP slope (*γ*_1_) and the quadratic curvature of the CM trajectory (*β*_2_), thresholded at *p* < 0.05. **(a)** For the *ξ*-process amplitude, we appreciate a widespread age-related decrease in ZIP (i.e., *γ*_1_ < 0) on the cortical surface, suggesting that the aperiodic component becomes increasingly detectable with age in all the regions. The CM exhibits strongly concave (inverted-U) trajectories globally, suggesting a rise in **A*ξ*** during early adulthood, followed by a decline with aging. **(b)** For the *α* -process amplitude, ZIP increases in frontal regions and decreases in occipital regions, revealing an anterior–posterior effective gradient in *α* localization across the lifespan. CM trajectories are significantly concave in posterior regions, consistent with the well-documented decline of **A*α*** in later life following a midlife peak. **(c)** For Peak Alpha Frequency (PAF), ZIP increases frontally and decreases posteriorly, paralleling the **A*α*** findings. Significant concave CM trajectories are predominantly confined to occipital regions, supporting a lifespan-related slowing of *α* frequency centered in posterior cortical areas.**(d)** Cortical average lifespan trajectories for each spectral feature (**A*ξ*, A*α***, and **F*α***) reveal robust inverted-U patterns with highly significant quadratic terms (*p* = 9.25 × 10^− 35^, 7.70 × 10^− 11^, and 6.46 × 10^−56^, respectively), indicating shared nonlinear developmental and aging dynamics across components. These results suggest that the cortical localization of the spectral process correlates with significant inverted U-shape developmental trajectories across the lifespan and an increase in localization.

**A*α*** [Fig. 5(b)] displays a distinct spatial gradient: ZIP increases in frontal regions and decreases in occipital areas, indicating that *α*-rhythms become less detectable anteriorly with age while remaining relatively stable posteriorly. CM trajectories for **A*α*** are also significantly concave in posterior regions, reflecting a peak in early-to-mid adulthood followed by an age-related decline in regions associated with *α* generation. For PAF or **F*α*** [Fig. 5(c)], we observe a similar ZIP gradient from occipital to frontal areas. The probability of zero inflation increases anteriorly, particularly in older individuals, suggesting that *α* peaks become less reliably detectable in the frontal cortex with age. Significant concave CM trajectories for PAF are localized to posterior areas.

The cortical average lifespan trajectories for each spectral feature are shown in [Fig. 5(d)]. All three components—**A*ξ*, A*α***, and PAF—exhibit robust inverted U-shaped trends, with highly significant quadratic curvature (*p* = 9.25 × 10^−35^ for **A*ξ***, *p* = 7.70 × 10^−11^ for **A*α***, and *p* = 6.46 × 10^−56^ for PAF).

These results are significant for several reasons. First, they confirm, at high spatial resolution and in a large cohort, the canonical inverted U-shaped lifespan trajectory of PAF in source space, as previously described in sensor-level studies [36,42–44]. Second, SC and spectral localization correlate with significant inverted U-shaped developmental trajectories, revealing that these patterns are not spatially uniform on the cortical surface. A posterior-to-anterior gradient of ZIP shapes the cortical distribution of detectable *α* activity, modulating the developmental curve, particularly in frontal regions.

### Estimated Conduction Delays are negatively correlated with the Peak Alpha Frequency and with independently reported myelin data

Conduction delays depend on axon diameter and myelin thickness, both of which govern the speed of action potential propagation. Myelination is the process by which lipid-rich sheaths wrap axons (myelin thickness increases). This process produces an increase in the speed of transmission of an action potential. As dMRI studies show, myelin expression is not a static structure established early in development; rather, it changes throughout the Lifespan. Myelin thickness increases from childhood to adulthood, reaching a plateau before starting to decrease with maturation [Fig. 6 (b)]. Consequently, myelin expression follows an inverted U-shaped trajectory across the Lifespan, modulating conduction velocities and neural dynamics [46]. According to the Rushton model [47], myelin thickness (*d*_*m*_) is inversely proportional to the square of the average conduction delay (⟨*τ*⟩):

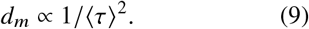

**Figure 6.**
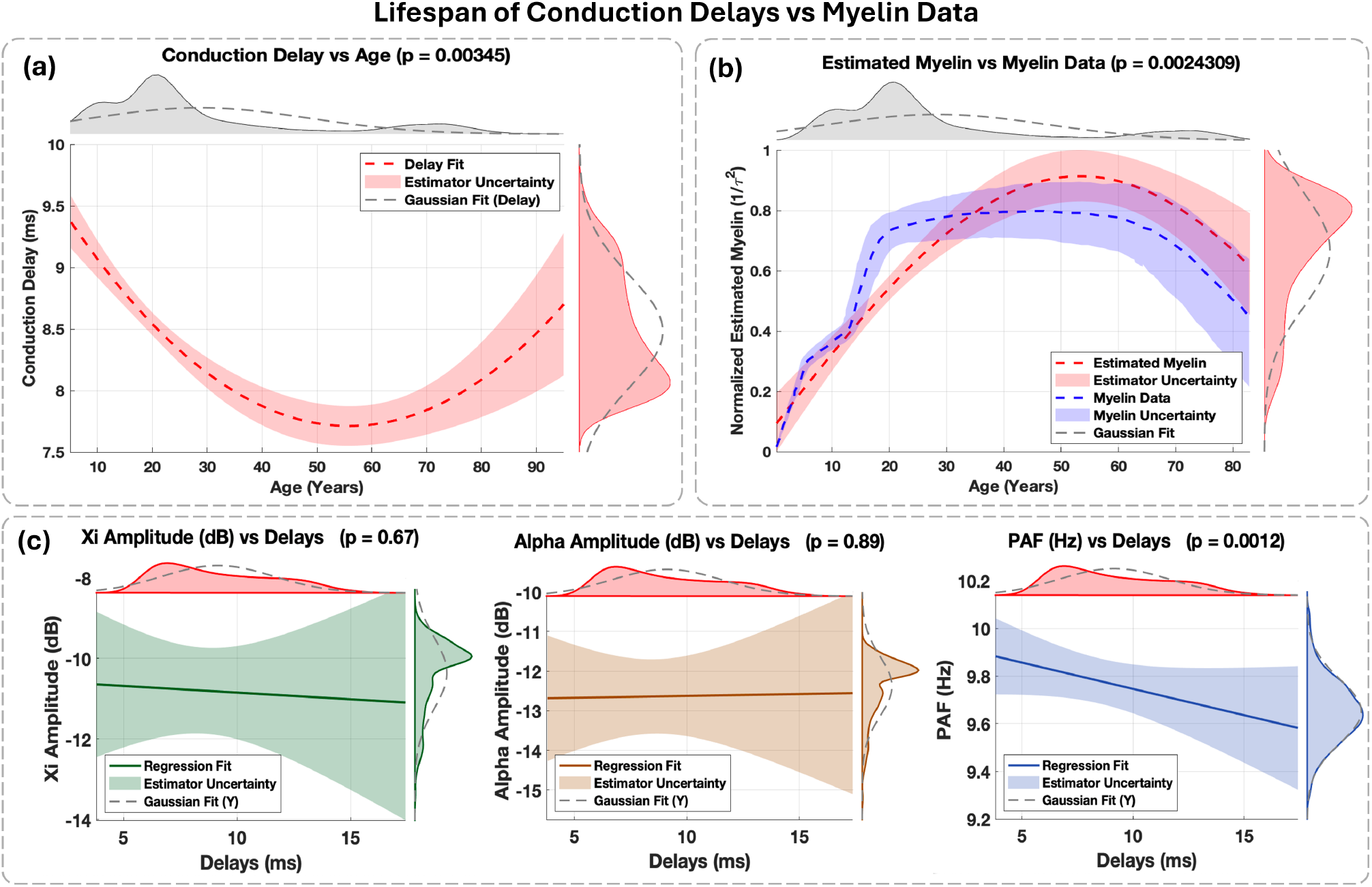
Lifespan of Conduction Delays and their relationship to Myelin and Spectral EEG Features. **(a)** Developmental trajectory of conduction delays estimated using robust quadratic regression on subject-level delays inferred from the *ξ* − *α* NET model in the HarMNqEEG dataset (*N* = 1965). The regression reveals a significant U-shaped relationship with age (*p* = 0.00345 for the quadrature), with a minimum occurring around early-to-mid adulthood. **(b)** Estimated delays are mapped to normalized cortical myelin content, revealing an inverted U-shaped trajectory that peaks in midlife (*p* = 0.00243 for the quadratic term). Overlaid in blue is an independent dataset of age-matched myelin measurements. The two curves show strong qualitative and quantitative agreement, both in shape and peak location (approximately 40-50 years), suggesting that the inferred delays are physiologically consistent with known patterns of myelination. **(c)** Robust linear regressions between conduction delays and EEG spectral features estimated from *ξ* − *α* NET: **A*ξ*** (*p* = 0.67), **A*α*** (*p* = 0.89), and **F*α*** (PAF, *p* = 0.0012). Only PAF exhibits a significant negative association with conduction delay, indicating that higher conduction speeds are associated with higher PAF values, in line with biophysical theories linking conduction velocity and oscillatory frequency.

Hence, the Rushton model and the trajectory of myelin thickness obtained using dMRI suggest that conduction delays should follow a U-shaped trajectory across the Lifespan. Nevertheless, these results have never been stated from EEG using an SCM. To obtain the trajectory of conduction delays across the Lifespan, our starting point is the F-TRACT atlas, which provides the matrix representing the average conduction delays for the age group > 15, as provided by Lemarechal et al. [35] [Fig. 2(b)]. This data informs the generative model of the *ξ* − *α*NET [Fig. 2(c)].

Specifically, subject-specific conduction delays ***τ*** ^( *j*)^ are estimated as:

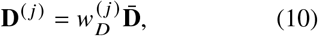

where 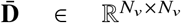 is the baseline conduction-delay matrix estimated by the F-TRACT group, and 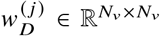 is a weight that modulates 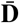 to reflect specific factors for subject *j* as they age. The weight 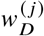 is then constrained within a reasonable search space for its estimation by the following condition:

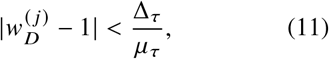

where Δ_*τ*_ = 5.5 ms and *μ*_*τ*_ = 9.5 ms denote the Median Absolute Deviation and the Median of the conduction delays, respectively, as reported by Lemarechal et al. [35]. The Bayesian Model Inversion of the *ξ* − *α*NET on the complete HarMNqEEG dataset allows us to learn 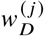 for each subject in the dataset. The average conduction delay ⟨**D**^( *j* )^ ⟩ across all neural tracts is then computed as:

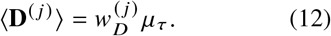

On these estimations, we perform a robust quadratic regression with age (*a* _*j*_ ) as the descriptor and the average conduction delay ( ⟨**D**^( *j* )^ ⟩) as responses. As shown in [Fig. 6(a)], the conduction delays follow a significant U-shaped trajectory across the Lifespan (*p* = 0.00345 for the quadratic term). Conduction delays decrease from the early stage of development to middle age, reaching a minimum around age 50, and increase again in later years. We then in [Fig. 6(b)], map. The estimated average conduction delays into normalized myelin proportion, which from the Rushton model is inversely proportional to the square of conduction delay, and over this estimated myelin proportion, conduct as before a robust quadratic regression with age as descriptor. This new estimated myelin proportion is compared to the developmental trajectory independently reported of myelination by De-Faria et al. [46]. As we can depict in [Fig. 6(b)], both the model-derived and empirical myelin curves follow a qualitatively and quantitatively similar inverted U-shaped pattern (*p* = 0.00243 for the model-derived quadratic fit), peaking between ages 40–50. These results suggest that the *ξ*-*α*NET model on rsEEG allows for estimating conduction delays that are physiologically plausible.

Importantly, regression of EEG spectral parameters against conduction delays revealed a significant negative correlation between PAF; **F*α***) and delay (p = 0.0012; [Fig. 6(c)]). In contrast, neither **A*ξ*** (p = 0.67) nor **A*α*** (p = 0.89) exhibited significant relationships. These suggest that faster conduction, which reflects higher myelin content, tends to show higher PAF values. Since PAF is the frequency of the dominant spectral peak of the *α*-rhythm, this finding supports the hypothesis that age-related changes in myelination influence the PAF trajectory, linking structural and functional maturation processes in the brain. This observation is consistent with earlier work by Valdés-Hernández et al. [48], who demonstrated a significant correlation between diffusion tensor imaging-derived fractional anisotropy (FA) and PAF on the Cuban Human Brain Mapping Project (N = 397) data. Furthermore, from the theoretical point of view, this result aligns with predictions from thalamocortical modeling studies by Robinson et al. [24,25], who proposed that delays within thalamocortical circuits modulate the emergence of damped *α* oscillatory modes.

These results not only confirm previous empirical knowledge but also represent the first time that rsEEG is used to estimate conduction delays that are compatible with the physiological nature of developmental changes on the basis of higher statistical power (HarMNqEEG, N=1965). Together, our results suggest that PAF is a sensitive marker of white matter integrity across the Lifespan and highlight the value of *ξ*-*α*NET in unraveling the interplay between structure and function in the human brain.

## Conclusion and Discussion

We introduce *ξ*-*α*NET, a generative model of EEG cortical activity that models each spectral component as a sparse, structurally constrained network of independent Hida-Matérn processes in the time domain. By integrating the connectome (derived from dMRI) and conduction delays (estimated from CCEP) [34,35,49], the model captures the spatiotemporal correla-tions between cortical generators and generates Lorentzian spectral profiles consistent with the seminal work on SCM by Pascual et al. [2,10,48, 50]. This formalism unifies spectral decomposition of both cortical activity and effective connectivity, addressing key limitations of existing SCM, which often neglect the role of structural priors on spectral estimates [6,9,10].

Using Bayesian inversion, *ξ*-*α*NET simultaneously estimates source spectral parameters and conduction delays from empirical data. Applied to the HarMNqEEG dataset (N = 1,965), this enabled the largest study to date linking rsEEG spectral dynamics and effective connectivity at the sources. The aperiodic component shows a widespread cortical distribution with significant inverted U-shape developmental trajectories alongside an age-related decline in zero inflation. Its effective connectivity patterns also revealed a widespread network. While in contrast, the *α*-rhythm is localized in the posterior areas, displaying two distinct specialized effective networks, a sensorimotor and parieto-occipital network. At the early stage of development, these specialized networks are disconnected, and the connection between them increases with age. On the other hand, both the amplitude and PAF displayed significant inverted U-shaped trajectories, with a pronounced posterior-to-frontal gradient in zero activations. These results are significant for two reasons. First, they replicate earlier probability atlases reported by previous studies based on smaller datasets [2,3,6,10,14,43]. Here, we confirm these topographical patterns with higher statistical power and spatial resolution. Second, by leveraging SGC, we independently delineate the effective connectivity networks of each spectral process for the first time, revealing marked differences in their causal architectures.

*ξ*-*α*NET also allows us to estimate conduction delays that closely mirrored independent myelin concentration profiles. These delay es-timations have a significant negative correlation with the PAF, establishing a direct link between electrophysiological properties and underlying structural maturation. All of these results align with both previous theoretical predictions from thalamocortical models by Robinson and prior statistical analyses of smaller datasets by Valdes, which reported significant correlations between fractional anisotropy and PAF [24,25,48].

Our work highlights the importance of modeling cortical activity using damped fractional stochastic differential equations with resonance, particularly Hida-Matérn processes, to capture generalized Lorentzian spectral profiles and provide physiologically meaningful interpretations of cortical dynamics. This implies that not only the aperiodic component but also all other SC can be understood as arising from fractional stochastic processes. From these, we conclude that the *ξ*-*α*NET framework is well-suited for large-scale normative studies of EEG data in the sources. Beyond that, it offers a low-dimensional representation of cortical activity to define an embedding for training large language models in both classification and prediction tasks. It holds promise for subject-specific neural field simulations, thereby supporting the development of personalized models of brain dynamics.

Despite its strengths, *ξ*-*α*NET framework has several limitations. Its simplification of thalamocortical activity into a linear MVAR model with colored Hida-Matérn innovations, while computationally efficient and accurate in some aspects, does not fully capture the complex excitatory-inhibitory balance that shapes network dynamics. The model also assumes that linear causal relationships between oscillatory activity drive network interactions, neglecting feedback mechanisms and bidirectional influences essential for capturing the true complexity of neural dynamics, which often exhibit nonlinearities, chaotic behaviors, attractors, and state-dependent transitions. Introducing nonlinear activation functions would allow us to capture these dynamic properties realistically. In this case, this is not done for simplicity. We use the Dirac delta distribution to model conduction delays may oversimplify the actual delay structures, which likely may require the use of more flexible distributions, such as the exponential family [28]. Although we believe the model can be applied to MEG signals, for which only the use of a different lead field will be required, we have not yet assessed the performance of the model on this type of data. This remains an open area for future investigation.

Moreover, the current focus on only two SC, the aperiodic (*ξ*) process and the *α* peak—limits its ability to capture other important rhythms such as theta, beta, and gamma. Incorporating these other SC as Hida-Matérn processes is important for modeling the full range of cortical dynamics. Finally, the model assumes stationarity, limiting its applicability to transient neural dynamics [7]. Addressing these limitations, including the introduction of nonlinear dynamics, will be the subject of future research aimed at enhancing *ξ*-*α*NET’s flexibility and accuracy in representing the full complexity of neural oscillations.

## Materials and Methods

### Resources and Code

All numerical computations and simulations were performed on a high-performance server equipped with 52 CPUs and 256 GB of RAM. The *ξ* − *α*NET parameter estimation pipeline, along with a visual interface for analysis, visualization, regressions, and documentation, is available on GitHub Xi-AlphaNET.

### Data

Two types of data were used: structural data for informing the *ξ* − *α*NET model and empirical data for analysis. For the structural data, we used the anatomical connectivity atlas 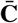 from Rosen et al. [34], which was derived from dMRI to estimate axonal fibers linking cortical areas. These estimates were calibrated using histological measurements of callosal fiber density and are available via the HCP-MMP1parcellation [21]. Axonal conduction delays atlas 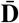 were obtained from the F-TRACTS consortium, based on data from 780 epilepsy patients [35]. These delays were preprocessed using Gaussian process regression and integrated with neurotract length because the original atlas contains missing data **L** [34] (see Supplementary Materials J). Both anatomical and conduction delay data were mapped from ROI space to voxel space. The head model, source model, and lead field matrix (**K**) were derived using the Ciftistorm pipeline, applying the HCP-MMP1parcellation [51].

Empirical data were sourced from the HarM-NqEEG dataset, which includes scalp rsEEG cross-spectral tensors from 1,965 subjects aged 5-100 years across nine countries and multiple recording devices. Each tensor has dimensions *N*_*c*_ × *N*_*c*_ × *N*_*ω*_, where *N*_*c*_ = 19 EEG channels and *N*_*ω*_ = 47 frequency bins. Preprocessing steps, as detailed in [36], included applying an aver-age reference, removing the global scaling factor, and ensuring the positive definiteness of the cross-spectral matrices (Supplementary Materials K & L). Visualization outputs were generated using the functions included in the BC-VARETA toolbox [17].

### Processed Data

The spectral and structural parameters estimated using the *ξ*–*α*NET model from the HarMN-qEEG dataset are available at the following location: Processed Data. The repository (∼ 40 GB) contains all estimated parameters, including the source power spectrum, cross-spectral matrices in source space, cortical activation maps, connectivity, conduction delay matrices, and the preprocessed structural data required to run the *ξ*–*α*NET model. For convenient visualization and validation, users are encouraged to use the Xi-AlphaNET app.

## AUTHOR CONTRIBUTIONS

R.G.R., L.M., and P.A.V.S. conceived and designed the study. R.G.R., P.A.V.S., and A.A.G. developed the Xi-AlphaNET software and visualization interface. R.G.R. conducted the formal analysis and wrote the original draft. L.M., C.L., P.X., D.Y., V.J., and M.L.B.V. contributed substantially to the interpretation and discussion of the results. R.G.R., Y.W., Y.J., and A.A.G. prepared visualizations. M.L.B.V., C.L., P.X., D.Y., V.J., and L.M. contributed to the manuscript review and editing. P.A.V.S. and L.M. supervised the project. P.A.V.S., L.M., and C.L. acquired funding for the research.

